# Case studies and mathematical models of ecological speciation. V. Adaptive divergence of whitefish in Fennoscandia

**DOI:** 10.1101/725051

**Authors:** Xavier Thibert-Plante, Kim Præbel, Kjartan Østbye, Kimmo K. Kahilainen, Per-Arne Amundsen, Sergey Gavrilets

## Abstract

Modern speciation theory has greatly benefited from a variety of simple mathematical models focusing on the conditions and patterns of speciation and diversification in the presence of gene flow. Unfortunately the application of general theoretical concepts and tools to specific ecological systems remains a challenge. Here we apply modeling tools to better understand adaptive divergence of whitefish during the postglacial period in lakes of northern Fennoscandia. These lakes harbor up to three different morphs associated with the three major lake habitats: littoral, pelagic, and profundal. Using large-scale individual-based simulations, we aim to identify factors required for in situ emergence of the pelagic and profundal morphs in lakes initially colonized by the littoral morph. The importance of some of the factors we identify and study - sufficiently large levels of initial genetic variation, size- and habitat-specific mating, sufficiently large carrying capacity of the new niche - is already well recognized. In addition, our model also points to two other factors that have been largely disregarded in theoretical studies: fitness-dependent dispersal and strong predator pressure in the ancestral niche coupled with the lack of it in the new niche(s). We use our theoretical results to speculate about the process of diversification of whitefish in Fennoscandia and to identify potentially profitable directions for future empirical research.

## Introduction

The diversity of species on Earth continues to provide inspiration for scientists studying speciation and the origins and maintenance of biodiversity. What makes these processes extremely complex and difficult to understand is that different evolutionary and ecological factors controlling their dynamics act simultaneously and often have opposing effects. The complexity of speciation processes implies that mathematical modeling can potentially play a very important role in uncovering its general rules and patterns. By now we have an impressive array of models and modeling techniques that shed light on the conditions, probability, waiting time to and duration of speciation, the degree of genetic and phenotypic divergence between the emerging species, and the way different resources (including space) are partitioned between the sister species. Models also explain the effects of different parameters and factors (such as the strength of selection, rates of mutation, recombination, migration, population size, number of loci, distribution of allelic effects, etc.) on the dynamics of speciation.^1–9^

Most of this work has focused on models of speciation aims for both generality and mathematical simplicity. These models are very useful and insightful in uncovering general rules and patterns of speciation, adaptive radiation, and biological diversification. However, their generality almost necessarily implies that these models are very difficult to apply to any particular systems and species studied by empirical biologists. Therefore it is very important to supplement general models of speciation with those tailored for specific biological cases.

Studying models tailored for particular case studies can be very useful from several perspectives.^9^ First, mathematical models emerging from these projects do lead to a better understanding of the evolutionary dynamics of the studied specific systems. Second, although the relevant models are case-specific, they contribute towards building the general theory of speciation, e.g. by supporting or undermining the generality of particular observations and patterns. Third, the process of building a mathematical model even for a particularly well-studied empirical system usually reveals the lack of biological understanding or crucial empirical data needed to make appropriate modeling assumptions or specify parameters. This can greatly stimulate further empirical work to remove these limitations.

By now a relatively small number of such models have been developed for some of the best studied systems. These include models aiming to capture the dynamics of non-allopatric speciation of cichlids in a lake^10–13^ and palms on an oceanic island,^14^ hybrid speciation in butterflies in Central America,^15^ ecomorph formation in marine snails in Sweden,^16^ pulmonate snails,^17^ and parallel adaptation in threespine stickleback.^18^

Here, we continue this work by focusing on a young and well-documented empirical system – the lacustrine European whitefish (*Coregonus lavaretus* (L.)) in Fennoscandian postglacial lakes. Ten to twenty thousand years ago Fennoscandia was covered by a thick ice sheet.^19^ The ice sheet retracted 8000 *−* 10000 years ago,^20, 21^ creating a landscape dominated by inter-connected lakes and rivers and thus providing the opportunity for postglacial immigration of cold-water adapted freshwater fish, such as whitefish. After the glacier melting and land uplifting, this region has a high density of lakes, with deep lakes having three major habitats (littoral, pelagic and profundal) for fishes. Almost all lakes in this region contain whitefish, but most lakes have only one littoral morph, some lakes two morphs (littoral and pelagic), and a few large and deep Fennoscandian lakes three morphs.^22–24^ The different morphs of whitefish show habitat specific patterns in their resource use, body size, gill raker number, and life-history traits.^22, 23^ Genetic evidence suggests a rapid divergence of these morphs from an ancestral monophyletic lineage, driven by natural selection on gill rakers and body size.^22, 25, 26^ Besides historical contingency, ecological opportunity mediated by interactions, such as resource competition and predation, may play an important role in diversification of whitefishes with regard to ecomorphology and life-history.^24, 27, 28^

What is remarkable about this system, compared to other postglacial lakes with pelagic and littoral morphs, is the presence of up to three distinct morphs in some lakes. This raises the question: could they have diverged within the lake, or have different morphs evolved in different lakes and came into contact later? Our goal here is to use individual-based simulations to better understand conditions for and various factors (abiotic and biotic) controlling the divergence of Fennoscandian whitefish into three principal habitats of subarctic lakes. Our particular focus will be on the effects of selection for local adaptation, gene flow, carrying capacity of the habitat, and predation.

## Methods

### Model

#### Environment

We consider finite sexual diploid populations inhabiting isolated lakes. Each lake has three different ecological niches (habitats) with its own whitefish population: littoral, pelagic, and profundal. Each lake can have up to four predator species feeding on whitefish: pike, perch, burbot, and brown trout (Table 1).^27, 29, 30^ Time is discrete, the unit of time is one year, and the generations are overlapping.

**Table 1:** Habitats and the maximum prey size for different whitefish predators.^27, 29, 30^

| Predator | Max prey size | Habitats |
| --- | --- | --- |
| Pike | 25cm | Littoral |
| Perch | 15cm | Littoral |
| Burbot | 20cm | Littoral, Profundal |
| Brown trout | 15cm | Littoral, Pelagic |

#### Individual whitefish

Each individual is characterized by three phenotypic traits: size *s*, the number of gill rakers *x*, and whether or not it is sexually mature. We discretize size *s* into four stages: 0, 1, 2 and 3, roughly corresponding to the size of 2cm, 15cm, 20cm, and 30cm in whitefish. Size stage 0 describes newly born individuals.

We assume that immature fish can grow while mature fish stop growing and invest all their available energy into reproduction. If an immature fish survives to the end of the year, it either matures, with a probability *m*_*s*_ depending on its current size *s*, or grows to the next size class *s* + 1, with probability 1 − *m*_*s*_. Newly born individuals can only grow but not mature (i.e., *m*_0_ = 0), whereas individuals reaching the largest size 3 always mature (i.e., *m*_3_ = 1). Note that in our model maturation always happens at the end of the year reflecting the need to accumulate energy for gonad development e.g.^31^ Figure 1 illustrates these assumptions.

**Figure 1:**
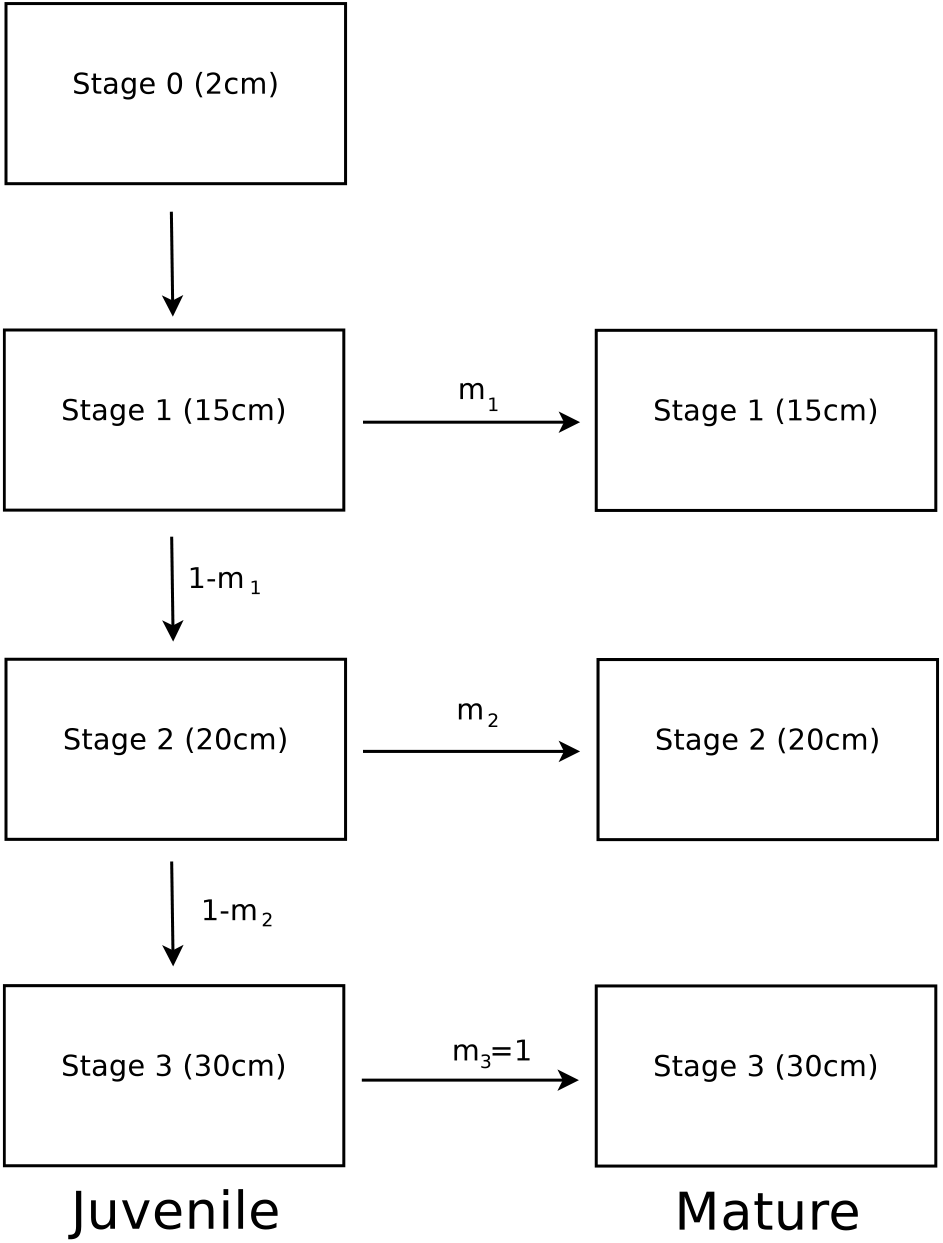
Life-history of a whitefish. The fish grows in size until it matures. The probabilities of maturation at stage 1 and 2 are *m*_1_ and *m*_2_, respectively. If a whitefish reaches stage 3, it definitely matures the next year and stops growing.

The studies on <u>Coregonus clupeaformis</u>^32, 33^ and salmonids^34^ show that maturation rate and the number of gill rakers are polygenic traits. Correspondingly, we assume that maturation rates *m*_1_ and *m*_2_ are additively controlled by *L* diallelic loci each. (In numerical simulations *L* = 4.) The corresponding allelic effects are scaled so that *m*_1_ and *m*_2_ stay between zero and one. The number of gill rakers *x* is also additively controlled by *L* loci but each locus has multiple discrete alleles. These alleles are subject to step-wise mutation and the effects are scaled so that trait *x* can take only integer values and mutation changes *x* only by plus and minus one. We assume that all genes are physically unlinked. Mutations happens with probability *µ* = 10^*−*5^ per gene per reproduction.

### Selection

We assume that the whitefish population is subject to density-dependent viability selection due to intraspecific competition, habitat-specific stabilizing selection on gill raker number, size-dependent mortality due to predation as well as fertility selection due to maturation rates and body size differences.

#### Condition

The gill raker number *x* controls the food available to fish in a given environment. We define the condition *ω*_*j*_(*x*) of a whitefish with *x* gill rakers in niche *j* as

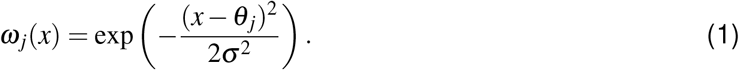

Here *θ*_*j*_ is the optimum gill rakers number in niche *j* and *σ* ^2^ is a parameter measuring the strength of stabilizing selection on *x* towards this optimum. For whitefish, we set *θ* at 26, 36, and 18 for the littoral, pelagic, and profundal habitats, respectively. These values are representative for well-established morphs and are close to those observed in whitefish morphs LSR, DR, and SSR discussed above.^24, 26^

#### Density-dependent selection

We assume that the population is subject to density-dependent mortality with the survival rate of individuals with *x* gill rakers and of size *s* in niche *j* being

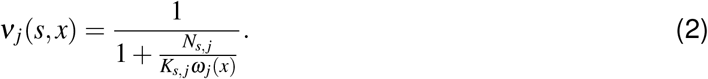

Here *N*_*s, j*_ is the total number of individuals of size *s* in niche *j*, and constant parameter *K*_*s, j*_ is the population size at which a population of perfectly adapted individuals (i.e., individuals with perfect condition *ω*_*j*_= 1) would have a survival rate of 0.5 (see the Appendix). We will interpret parameters *K*_*s, j*_ as measures of carrying capacity of the corresponding sizes in the corresponding niches. The larger the number of competitors *N*_*s, j*_ and the worse is the individual condition *ω*_*j*_, the smaller is the survival rate *ν*_*j*_. Our formulation follows the Beverton-Holt model^35^ and implies that density-dependent competition occurs only between individuals of the same size in the same ecological niche. The latter assumption reflects that fact that resource partitioning among whitefish morphs is very strong e.g.^23, 36^ and that intra-morph resource competition is much higher and the most pronounced between individuals of nearest age cohorts and size classes using same resources at the same habitats.^37, 38^

#### Predation

Maximum gape of a predator restricts the sizes and shapes of prey that can be eaten. We assume that each predator species in our system is characterized by a maximum prey size *g*_*k*_. To describe predation, we posit that predator *k* removes a random proportion *π*_*k*_ of surviving whitefish with sizes smaller than or equal to its gape size *g*_*k*_. Values *π*_*k*_ are parameters in our model. When whitefish are bigger than the maximum prey size of a predator, they are not predated upon. The maximum prey size of the four predators for whitefish are approximated in Table 1. Note that each predator is found in only a subset of habitats.

### Dispersal

Individuals can change their ecological niches within the lake. We assume that, each year, after reproduction each fish chooses the ecological niche to go to with probabilities proportional to their overall survival in each niche (accounting for both density-dependent selection and predation). One interpretation of this assumption is that each fish samples different niches before deciding on the one to stay. Fitness-dependent migration has been used in a number of earlier ecological models^39–42^ but has apparently been neglected in speciation modeling. Note that after each dispersal event, the niche population sizes change, so niche “attractiveness” can change as well. In numerical implementation of the model, to avoid a bias due to a sequence of events during the dispersal, individuals are chosen at random (without replacement, so that individuals move, at most, once per time step) to make a dispersal move.

### Reproduction

We assume that mating is assortative by the size and the niche. [This modeling choice can also accommodate a scenario where fish do not mate within their physical niches but rather mate at a niche- and size-specific mating ground or mating time.] Each surviving female chooses a mate at random from an appropriate set of males. She then produces a random number of offspring sampled from a Poisson distribution with size-specific means *b*_1_*, b*_2_ and *b*_3_. All offspring are born in the same niche as their parents.

### Life cycle

We assume there is a yearly sequence of events in the life of every fish. It starts with random death of eggs which is followed by density-dependent survival which is followed by predation mortality. The surviving fish grow or reproduce. Then all fish, except the newborn, have the opportunity to disperse among the niches within the lake, before this cycle starts again.

## Numerical simulations

### Scenarios

We simulated two scenarios of diversification differing in initial conditions. The Colonization-L scenario reflects the current view of the post-glacial colonization of lakes by large littoral fishes.^24^ In this scenario, the initial population of size 3,000 eggs is introduced in the littoral niche to which it is adapted. Its modal trait values are: *x* = 26, *m*_1_ = 0, and *m*_2_ = 1, that is, fish grows to the “large” stage 2 only. The Colonization-G scenario is similar as Colonization-L scenario, but all initial colonizers are assumed to grow bigger, i.e. reach the “giant” stage three (*x* = 26, *m*_1_ = 0, and *m*_2_ = 0).

In both scenarios, the initial population harbors genetic and phenotypic variation. The standard deviation of the gill raker number is five (which is close to values observed in natural populations). In the genes controlling the probabilities of maturation, one allele has frequency 95% and another has frequency of 5%. After introduction, the population then evolves for 10, 000 years. We then evaluate if and how many different morphs emerge and are maintained in the lake.

### Parameter values

In numerical simulations, besides the initial conditions, we also varied the predation intensity *π*. Specifically, for each of the four predators we set *π* at 0%, 30%, and 50%.

In our model, fertility parameters {*b*_1_, *b*_2_, *b*_3_} are set to {4, 16, 64} respectively, following an exponential relationship with length.^43, 44^ Note that these numbers should be interpreted not as an actual number of eggs produced by a fish but rather as a number of offspring surviving to the moment when they are subject to selection.

To estimate carrying capacities *K*_*s, j*_ (used in eq. (2)) we used available data assuming that extant fish are adapted to their environment, i.e. that they have the optimal number of gill rakers and reach maturation at a proper size (*m*_1_ = *m*_2_ = 0 for the littoral niche, *m*_1_ = 1 for the pelagic niche, and *m*_1_ = 0, *m*_2_ = 1 for the profundal niche). We also assumed that extant fish in three niches are reproductively isolated and that fertility is a function of the female size.^43, 44^

We set the equilibrium densities of the largest morph in each niche to 1,000, 6,000, and 150 individuals for the littoral, pelagic, and profundal niche, respectively. These values are proportional to the observed densities in each niche.^45, 46^

From these values and empirical data, we estimated the corresponding equilibrium densities for other morphs (Table 2) and the values of corresponding carrying capacity parameters *K*_*s, j*_ (Table S1 in the Appendix).

**Table 2:** Equilibrium population densities *N*_*s, j*_ for fish of different sizes in different habitats in the baseline model.

| Stage | Littoral | Pelagic | Profundal |
| --- | --- | --- | --- |
| 0 (2cm) | 15125 | 5517 | 656 |
| 1 (15cm) | 3025 | 6000 | 66 |
| 2 (20cm) | 756 | 0 | 150 |
| 3 (30cm) | 1000 | 0 | 0 |

The carrying capacities of different niches vary among different lakes. A particularly important factor affecting the likelihood of diversification is the carrying capacity of the profundal niche which can vary because of the size of the lake, its shape, and the nutrient deposition rate. In this study we used three different carrying capacities for the profundal niche resulting in equilibrium densities equal to one, two, and four times the values given in Table 2. We kept carrying capacities of the other two niches constant.

Overall, the results we present here explore 2 × 3^4^ *×* 3 = 486 different combinations of parameters, assuming assortative mating and fitness-dependent dispersal, for each of which we did 10 independent stochastic runs. We also performed a similar investigation of four related sets of models: 1) with random (rather than size-assortative) mating and random (rather than fitness-dependent) dispersal, 2) with size-assortative mating and random dispersal, and 3) with random mating and fitness-dependent dispersal. We also investigated our model with 4) smaller initial standard deviations in gill raker number and lower variation in the maturation loci. We never observed the emergence of the profundal morph in these models so we do not present the corresponding results.

### Evolutionary outcomes

To interpret our numerical results, we say that a given niche has been successfully colonized if in the last year the population of newborn both 1) has gill rakers number adapted to their niche and 2) they mostly grow to a size which characterizes the niche in real lakes.

The former assumption was formalized as the requirement that the average gill raker number *x* is close to the optimum value, specifically: *θ*_*j*_ − *σ* ≤ *x* ≤ *θ*_*j*_ + *σ*. (Recall that *σ* is a parameter measuring the strength of stabilizing selection.) The latter assumption was formalized as requirements on the evolved average maturation rates *m*_1_ and *m*_2_. Specifically,

- if 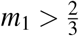 (i.e., fish mostly matures at size 1), then we say the fish is pelagic;
- if 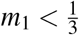,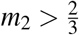 (i.e., fish mostly matures at size 2), then we say the fish is profundal;
- if 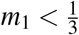, 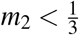 (i.e., fish mostly reaches size 3), then we say the fish is littoral.

## Results

Here we present our main results on the conditions under which each niche was colonized and by how many morphs. [Our results for the entire parameter space studied can be found online (https://bit.ly/2qCFR6P) (username: Scandinavia2, password: Evolution45! - the password protection will be removed when the paper is published). The online browser will be archived on DRYAD (DOI available later).]

When we look across the entire parameter space, the majority of lakes fall into four different compositions of morphs present: littoral morph only, pelagic morph only, littoral and pelagic morphs, and all three morphs present (Tables 3 and 4). In most case, the system reached a stochastic equilibrium state on the time scale of a few hundred generations (see the link above for some examples). This observation on the speed of diversification is comparable with earlier speciation models (e.g., gavrilets2004,gavriletsVose2005,gavriletsEtAl2007). Below we will break down the conditions under which each outcome was more likely to be observed.

**Table 3:** Number of simulations with different compositions of morphs present for different relative carrying capacities of the profundal niche in Colonization-L scenario.

| Relative carrying capacity | 1.0 | 2.0 | 4.0 |
| --- | --- | --- | --- |
| None | 2 | 2 | 1 |
| Monomorphic | 479 | 495 | 509 |
| Littoral | 439 | 477 | 501 |
| Pelagic | 39 | 17 | 8 |
| Profundal | 1 | 1 | 0 |
| Dimorphic | 329 | 313 | 295 |
| Littoral Pelagic | 329 | 313 | 279 |
| Profundal Littoral | 0 | 0 | 16 |
| Profundal Pelagic | 0 | 0 | 0 |
| Three morphs | 0 | 0 | 5 |

**Table 4:** Number of simulations with different compositions of morphs present for different relative carrying capacities of the profundal niche in Colonization-G scenario.

| Relative carrying capacity | 1.0 | 2.0 | 4.0 |
| --- | --- | --- | --- |
| None | 0 | 0 | 0 |
| Monomorphic | 425 | 484 | 520 |
| Littoral | 422 | 482 | 519 |
| Pelagic | 3 | 2 | 1 |
| Profundal | 0 | 0 | 0 |
| Dimorphic | 385 | 326 | 290 |
| Littoral Pelagic | 385 | 326 | 282 |
| Profundal Littoral | 0 | 0 | 8 |
| Profundal Pelagic | 0 | 0 | 0 |
| Three morphs | 0 | 0 | 0 |

### Only littoral morph (no diversification

Survival of the littoral morph with no other morphs emerging was the most common outcome in our simulations (Tables 3, 4 and Figures 2, 3). It typically required the presence of predation by trout. Lakes with only the littoral morph are common in Fennoscandia.^24^ Their abundance in the geologically younger region is one of the reasons that we hypothesize that the littoral morph is ancestral to Fennoscandia lakes.

**Figure 2:**
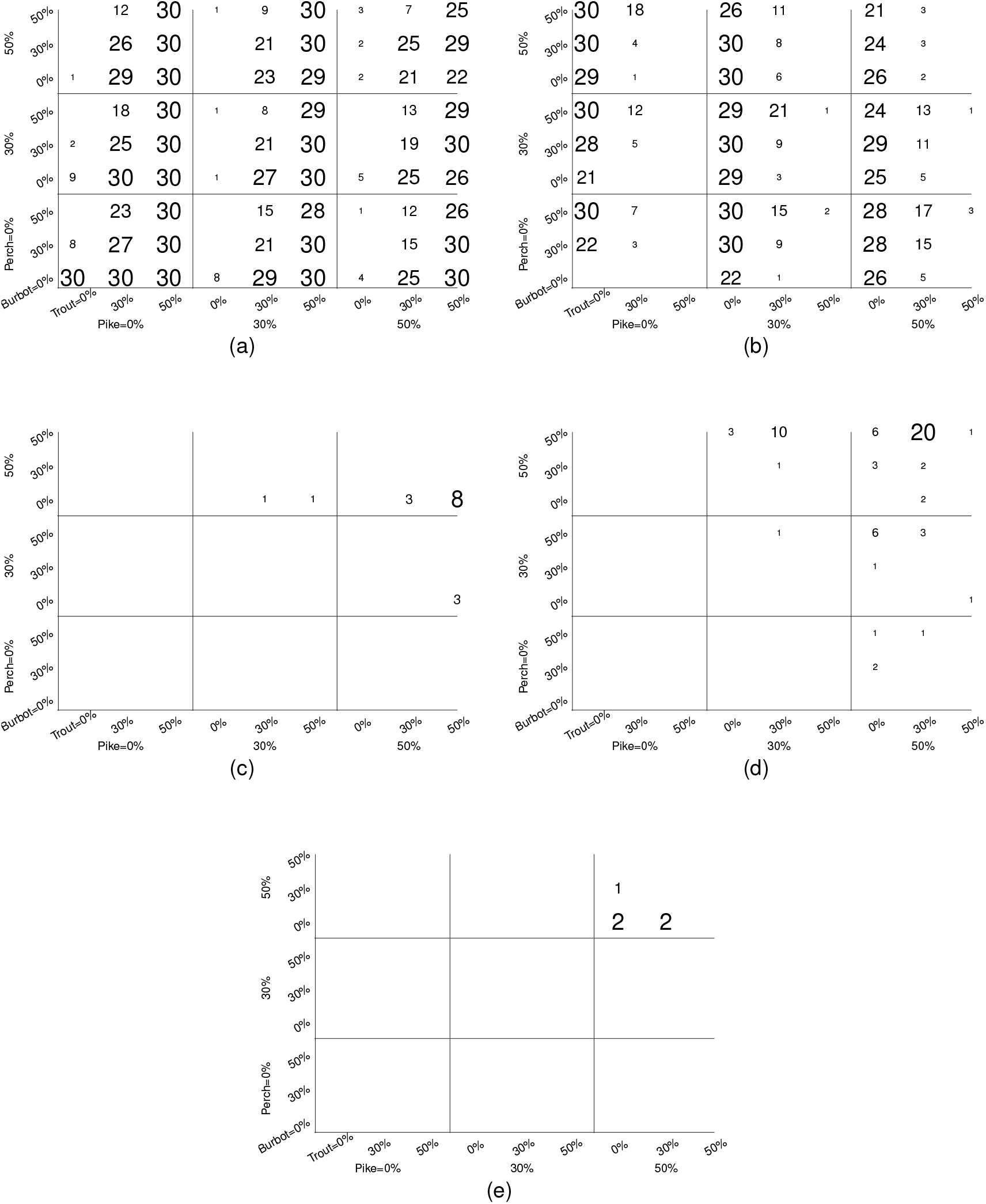
The frequencies of different outcomes in the Colonization-L scenario for different predation rates. 30 simulations for each parameter combination. The numbers are also reflected in the size of the font used. (a) Only littoral morphs. (b) Both littoral and pelagic morphs. (c) Both littoral and profundal morphs. (d) Only pelagic morph. (e) All three morphs.

**Figure 3:**
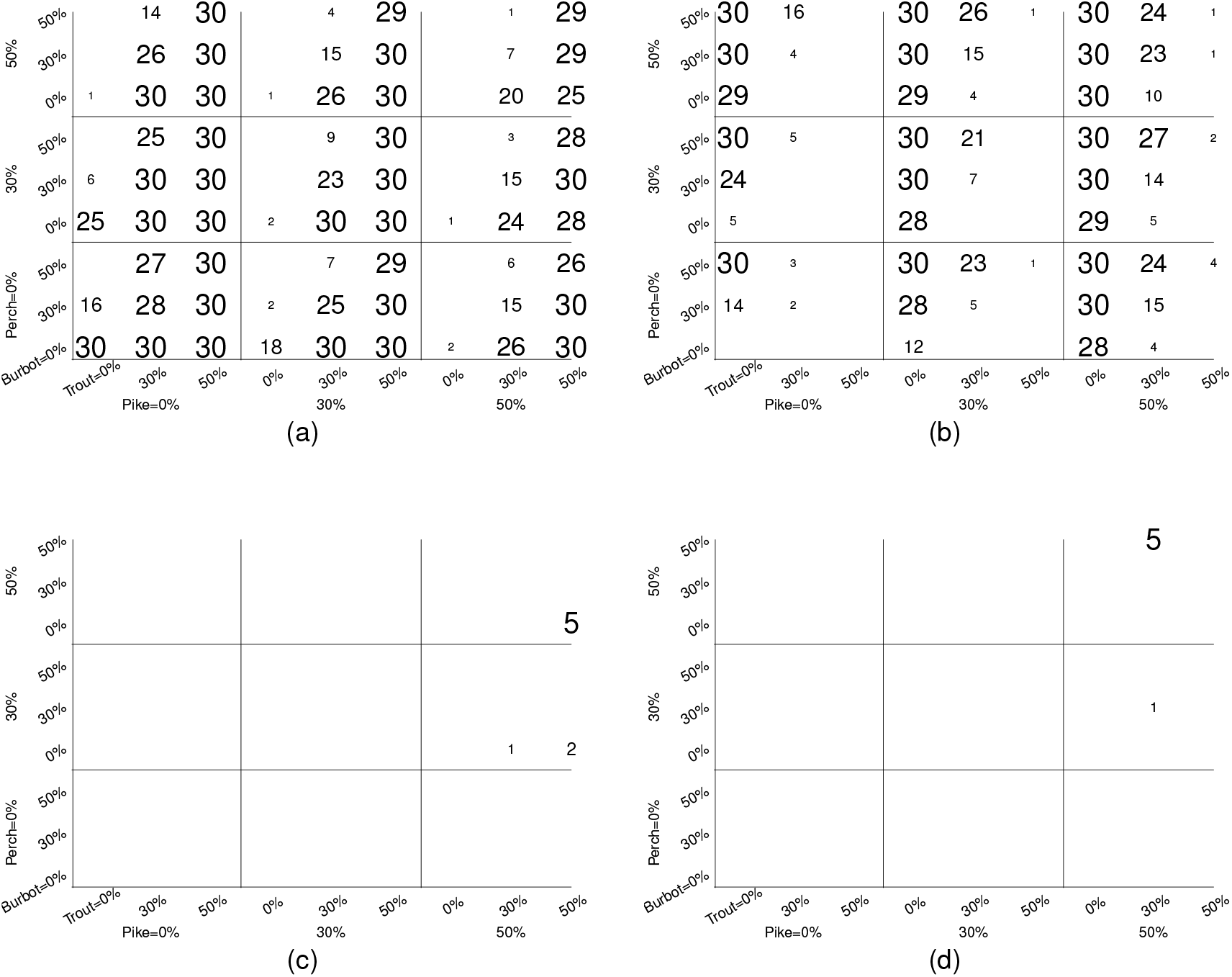
The frequencies of different outcomes in the Colonization-G scenario for different predation rates. 30 simulations for each parameter combination. The numbers are also reflected in the size of the font used. (a) Only littoral morphs. (b) Both littoral and pelagic morphs. (c) Both littoral and profundal morphs. (d) Only pelagic morph. No cases with all three morphs were observed.

### Two morphs

The emergence of the pelagic morph and the maintenance of the littoral morph was a frequent outcome in both colonization scenarios (about 40%). This outcome occurs more often when trout predation was absent or small ((Tables 3, 4 and Figures 2, 3)). There were also a few cases (24) where the divergence occurred toward the profundal morph, instead of the pelagic (Tables 3, 4 and Figures 2c, 3c). All these cases were observed when the profundal environment was large and no burbot predator was present. This outcome is usually not observed in Fenoscandia.

### Single pelagic morph

Although lakes with only the pelagic morph present are not observed in Fennoscandia,^24^ but they were observed in our simulations. This outcome implies the extinction of the littoral morph after it gives rise to the pelagic morph. This outcome was observed in both colonization scenarios under high predation by pike and perch but much more common in the Colonization-G scenario (Tables 3, 4 and Figures 2d, 3d).

### Three morphs

The cases with all three morphs present were very rare in our simulations - 5 cases only. They all happened in Colonization-L scenario under large carrying capacity for the profundal niche (Table 3) when predation from pike and perch was strong but that from trout and burbot was weak (Figure 2e). Lakes with all three morphs are rarely observed in Fennoscandia.^24^

### Failed adaptation

Four simulations resulted in populations with intermediate traits according to our metrics. These failed adaptations all involved high predation. Additionally, in one simulation with strong predation, the population went completely extinct.

Overall, both monomorphic and polymorphic lakes emerged from our simulations. The outcomes of Colonization-L and Colonization-G scenarios were similar, but Colonization-L scenario was more prone to diversification. The reason is that Colonization-L scenario started with the gill raker numbers typical for the littoral morph but with the size close to those for the pelagic and profundal morphs. This simplified the emergence of these two morphs. The lakes with all three niches colonized were observed but very rarely.

## Discussion

We used individual-based simulations to study the likelihood of within-lake ecological and morphological diversification in Scandinavian whitefish in the post-glacial time frame (about 10, 000 years). Our simulations show a common emergence of lakes where, in addition to the originally colonizing littoral morph, either the pelagic or profundal morphs have evolved and become established. The former outcome was much more frequent than the latter. We also observed the presence of lakes with all three morphs present, although very infrequently. However, this result fits with the observations from the wild which have revealed only a handful of trimorphic systems.

We started our simulations with a small founder population adapted to the littoral niche and no individuals adapted to the two other available niches: pelagic or profundal. Colonizing the pelagic niche required individuals to evolve increased gill raker number and smaller size. Colonizing the profundal niche required individuals to evolve decreased gill raker number and smaller size. The formation of new morphs typically happened on the time-scale of a few hundred generations. This is similar to other models with some initial genetic variation and strong selection.^7, 47, 48^

Speciation theory tells us though that the mere existence of “empty” ecological niches does not guarantee that they will be colonized and “filled” by locally adapted organisms, especially if there is a possibility of gene flow from the ancestral niche. Some additional factors must usually be in place to simplify survival in the new environment and reduce the effects of deleterious gene flow preventing local adaptation. Several such factors turned out to be very important in our model as well.

First, populations must have sufficiently large initial genetic variation. In our simulations presented above, we set the initial standard deviation of the gill rakers at 5. With a lower standard deviation (and, correspondingly, lower initial genetic variation), the state with both littoral and pelagic morphs was possible, but the profundal morph never emerged. Second, diversification required mating to be assortative with respect to both size and ecological niche. Third, dispersal had to be fitness-dependent. Both these assumptions result in reduced effects of gene flow from the ancestral niche. With random mating and/or dispersal, empty niches remained largely empty. Colonization of the profundal niche was mainly observed when its carrying capacity was the largest. Biologically, this describes lakes that are large and deep. Large carrying capacity simplified survival in the new niche by effectively reducing within-niche competition. These four assumptions alone were still not enough to ensure diversification. The fourth important factor was the elevated predation pressure in the littoral niche. Colonization of the pelagic niche largely required the absence of brown trout which is the only pelagic predator in model assumptions. Correspondingly, colonization of the profundal niche largely required the absence of burbot which is the only predator that can be found there. By starting to utilize the niche with no predators, the fish gets an immediate additional survival advantage which does not require evolving any genetic adaptations. Moreover, the absence of predators in a niche implies that fish does not have to grow large and thus could evolve earlier maturation. The latter in turn would lead to faster population growth.

The appearance of the pelagic morph in addition to the littoral one was much more common than that of the profundal morph, which is in accordance with findings from empirical studies.^24^ This happens because in our model, reflecting the situation in most lakes, the carrying capacity of the pelagic niche was much larger than that of the profundal niche making the pelagic niche much easier to colonize. Diversification was promoted if initial colonizers were littoral fish of “large” (Colonization-L scenario) rather than “giant” size (Colonization-G scenario), which happens because the size of the former is closer to those of the two other morphs (that are smaller than the littoral) and thus much smaller evolutionary change is required. However, body size is very plastic in whitefish and influenced by resource availability and strength of competition (e.g. amundsenEtAl2004, kahilainenEtAl2005). The Colonization-G scenario may nevertheless be relevant as large size likely is beneficial for migration. Following this scenario, “giant” colonizers may initially have experienced untapped resources and fast growth rates, but subsequently turned into the “large” category due to reduced growth rates from increasing fish density and intraspecific resource competition. The modeling results corroborate with a recent empirical study from northern Fennoscandian whitefish indicating that body size and number of gill rakers are both targeted by natural selection.^26^

Next, we discuss whether the factors identified in our model are present in natural populations of whitefish. Large genetic variation in colonizing individuals may have derived from the ancestral populations in the postglacial refugia, hybridization events during the loss of their pre-glacial habitats, and ongoing hybridization in contemporary times. Hybridization has an important role in historical and contemporary evolution of whitefish.^49, 50^ Whitefish is well-known for its high capacity for both fast forward and reverse speciation, which suggest a role of hybridization behind the high genetic variation harbored by the whitefish.^22, 25^ In the current modeling approach the minimum variation of five gill rakers of the ancestral morph was needed for the divergence to other morphs. Such variation of gill raker number is typical for generalist whitefish throughout much of the distribution suggesting ecological relevance of the model assumptions. However, the very few lakes with all three morphs in the simulation model overlooks the early individual specialization to all three habitats by the ancestral colonizing morph.^51^ Such step is very likely a crucial initial step towards full-fledged ecological speciation described as five model simulations leading to occurrence of all three morphs. While the proportion of simulations leading to three morphs is around 0.6%, it could be indeed comparable to natural conditions where this morph is present only at very few large lakes having wide profundal habitat coverage and many predators in littoral and pelagic niches.^27, 52^ Furthermore, frequent occurrence of pelagic and littoral morphs (36-40%) of simulations obviously refers to frequent divergence to habitats providing main energy sources (pelagic phytoplankton and littoral algae/macrophytes) to the lake food webs. In contrast, the profundal habitat is an unproductive habitat and dependent on energy inputs (settling materials) from the pelagic and littoral. These pelagic and littoral inputs vary among systems due to lake morphology and productivity and this complexity makes whitefish divergence to this habitat rare.^24, 52^

Size assortative mating is common in fish.^53–55^ Body size and size at sexual maturation of whitefish morphs are highly correlated with their resource use. Specifically, specialization to littoral benthic prey leads to large size and late maturation, pelagic prey to smallest size and fastest maturation, and finally profundal benthic prey to intermediate body size and late maturation.^22, 23, 37, 38^ Such differences in body size and maturation would be a logical step towards size assortative mating, if similar sized mature individuals among morphs are rarely present at the spawning sites. At least in the pelagic morph this seems likely as the very high mortality of small pelagic morph suggest that very few individuals are able to survive and grow to the size when other morphs mature.^37, 38^ Reproductive isolation may be further strengthened by temporal and spatial differences in time and place of spawning.^56, 57^ Such differences in spawning may arise via the habitat specific differences in temperature that regulates both prey resource availability and initiation of spawning.^31, 58^ In Fennoscandia, the littoral benthic resources are at the highest from early to mid-summer, pelagic zooplankton at mid to late summer and profundal benthic resources at late season.^31, 37, 38^ The littoral reaches the highest water temperature in summer, but cools down at the earliest time followed by the pelagic and profundal.^59^ Differences in realized water temperature among the whitefish morphs may be a major driver of divergence in Fennoscandian whitefish,^**?**^ as divergent temperature regimes alter the maturity status of all three morphs. There is also some evidence of spatial divergence of spawning sites among the morphs and in many salmonids such spawning site fidelity maintains population divergence.^60–62^ In our model, we assumed that individuals exhibit certain habitat preferences. In fish, those can take different form such as natal environment imprinting, condition dependent, density dependent, and predator avoidance.^63, 64^

In the model, the lack of pelagic (brown trout) and profundal (burbot) predation was a needed prerequisite for the raise of both pelagic and profundal whitefish morphs. Such conditions where strong predation occurs solely on littoral habitats of large lakes seems unlikely in contemporary conditions.^27, 29^ However, in the large lakes without polymorphic whitefish, brown trout and other predators (pike, burbot, perch) indeed used mostly littoral habitat.^52^ It is likely that the raise of pelagic whitefish morph induce brown trout shift to pelagic habitat use and piscivory.^30, 52^ Similar process could be present with regard to burbot, where divergence of profundal morph provides a new forage fish. However, burbot is a dark active predator that frequently use diel bank migration to feed on more abundant littoral prey resources.^65^ Thus, the empirical data suggest that modeling requirements for predation could be met in nature, but such conditions are rare.

Due to computational considerations, our model has some obvious limitations. For example, we assumed that fish laid a relatively small number of eggs. In reality, fish can produce a very large number of eggs; much larger than what we could simulate using an individual-based approach. We expect that with larger number of eggs, selection will be more efficient and divergence may occur faster than what is observed here (cf. gir10). The initial conditions were centered only on scenarios of “giant” and “large” colonizers, but it is very difficult to determine the actual body size of the ancestor starting to diverge. However, increasing intraspecific competition for resources is likely after the colonization of a new lake, which tend to shrink the body size supporting shift from giant to smaller body size. Predation by multiple species had a strong effect on whitefish divergence, but in the wild there are lake systems with whitefish morphs without main predators or very low amount of predation.^26^ However, the majority of lakes with three morphs have abundant predator populations and predation is likely to have strong influence on life-history divergence of prey. It is also likely that prey and predators co-evolve during the divergence process^66^ but individual-based modeling of multi-species evolutionary processes is inherently difficult. The process of building the model has also revealed important gaps in the available data on whitefish populations and their environments. In particular, having more precise estimates of the population sizes, predation rates, and fecundity of different morphs would increase the power of our model.

In our model, the body size was subject to direct selection by predation and also due to fertility and maturation rate differences. We did not consider explicitly body size adaptation to the ecological niche. However since body size correlates with gill raker number,^23, 36, 67^ our model partially captures this effect.

The sizes and depth of lakes in Fennoscandia vary a lot as well as their productivity with evident implications to species divergence.^24^ In our simulations, we considered only three major lake habitats and assumed pre-determined carrying capacities for each habitat. Lake morphometry and productivity in addition to de-glaciation history likely fine-tunes the divergence processes further,^24^ but these are far too complex scenarios to model conclusively. Further modelling effort is needed to understand vertical pelagic divergence documented for example to Alpine whitefish^68^ and North American ciscoes.^69^ Also, it is well established that a small number of loci of large effect are more conducive to speciation than a large number of loci with small effects.^7, 48, 70^ We have chosen a small number of loci to run our simulation to facilitate the process of adaptive radiation. With larger number of loci, we would not expect much diversification to be observed in our model. Given that diversification in whitefish does happen, we expect that the underlying traits are controlled by a small number of loci with large effects.

Overall, our modeling supported the possibility of divergence to three lake habitats during the postglacial time-frame, although such cases were rare and required a large profundal habitat without predators. Divergence to pelagic and littoral morphs was much more frequent and occurred with various levels of predation. The modeling effort indicated how little actually we know about reproductive isolation mechanisms as well as about habitat-specific carrying capacities and essential population dynamic parameters such as age-specific mortality in whitefish. Nevertheless, it would be interesting to apply our modeling approach to a true deep water species, such as Lake Baikal <u>Cottus</u> and various members of <u>Salvelinus</u>, in very large lakes using species-specific parameters.^71–73^ Recent empirical studies have highlighted the important role of deep water habitats which have been previously neglected in monitoring efforts.^74, 75^ In these respects, mathematical modeling could provide an efficient predictive tool for finding new lakes with three morphs, as well as identification of key factors contributing to their disappearance from the postglacial lakes.

## Acknowledgments

We thank Göran Englund, Jung koo Kang, Abisko Research Station, Polar Research grant, the Norwegian Research Council (grant no. 186320), Centre for Ecological and Evolutionary Synthesis (CEES), University of Oslo, the Office of Naval Research grant W911NF-17-1-0150, and the National Institute for Mathematical and Biological Synthesis supported through the NSF awards EF-0832858 and DBI-1300426, with additional support from The University of Tennessee. We have no conflict of interest to declare.

## Appendices

### Demographic equilibrium

To estimate the carrying capacity parameters *K*_*s, j*_ we proceeded in several steps. First, we estimated the birth rate for whitefish of different size using data in.^43, 44^ Second, using personal observations on the proportions of mature fish in each niche for trimorphic lakes we came up with realistic values of their equilibrium densities (*N*) at 1000, 6000 and 150 individuals, for littoral, pelagic and profundal morphs, respectively. These values serve as default values in our simulations.

**Figure S1:**
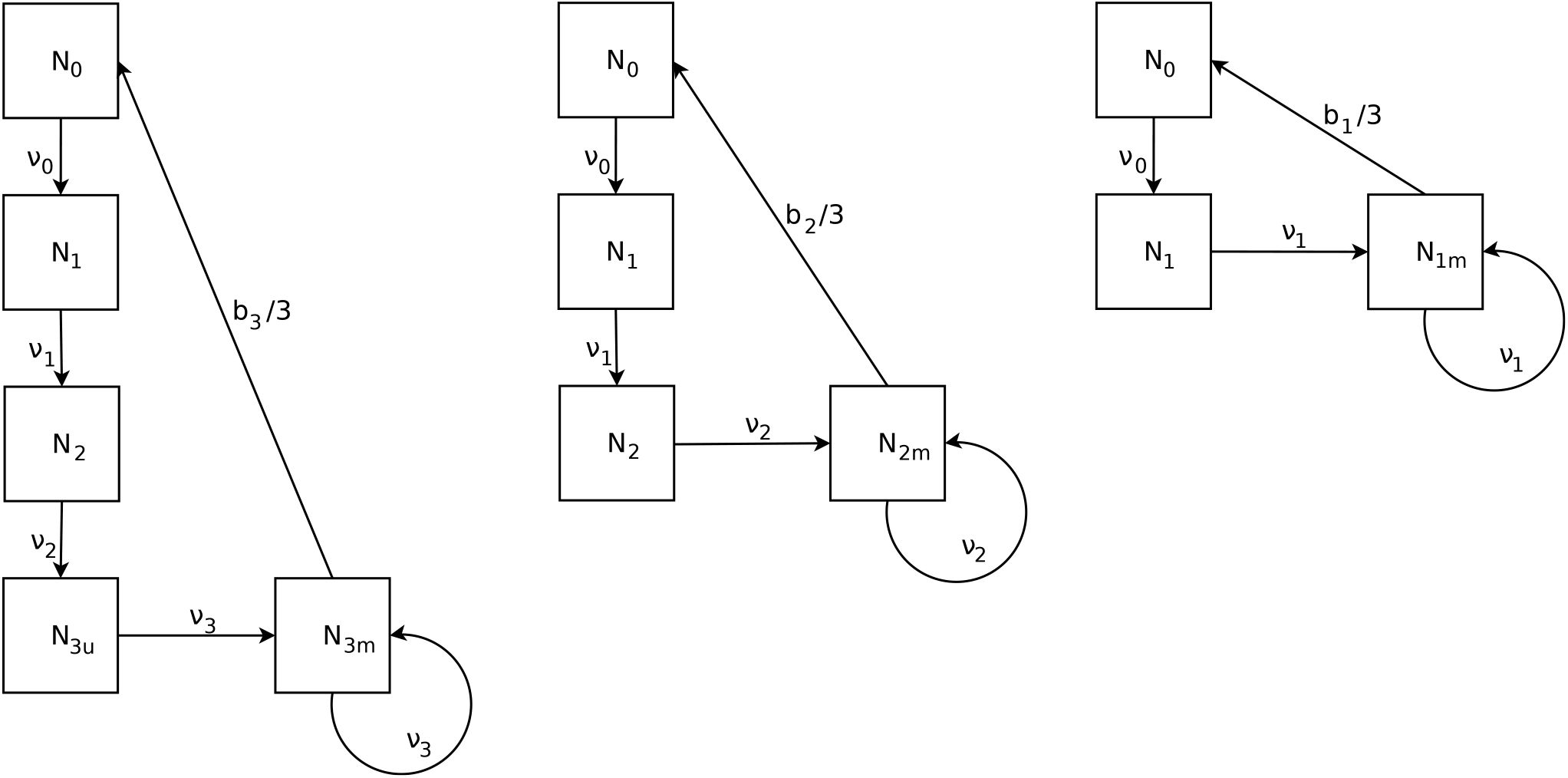
Demographic analysis of each population. *N*_*s*_, *v*_*s*_ and *b*_*s*_ are the population density, survival rate, and fertility at size *s*, respectively. Subscripts *u* and *m* specify immature and mature individuals.

Then, using the Leslie matrices approach (see Figure S1) and assuming perfect adaptation and no predation, we find that the equilibrium population densities *N*_*s*_ of fish of different size in different niches must satisfied the following equalities:

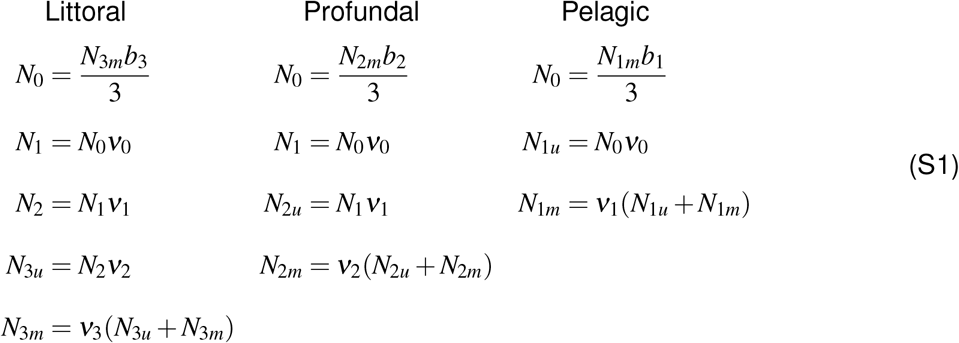

where *b*_*s*_ is the average number of offspring per female of size *s*, and subscripts *u* and *m* denote immature and mature individuals, respectively. Note that the division by three in all equations for *N*_0_ above is due to the 2:1 male:female sex ratio in whitefish (Author personal observations).

#### Littoral morph

Exact data needed to estimate all parameters of our model are difficult to come up with. As a result in some cases we have to take an educated guess. Using our unpublished data (Author unpublished), we know that one of the 2, 048 sampled littoral fish survived up to 26 years, thus living 23 years at stage three. This gives the yearly survival rate of *ν*_3_ ≈ 0.71. We then can find a relation between the mature and immature population sizes at stage three. With *N*_3_ = 1000, we have *N*_3*m*_ = 709 and *N*_3*u*_ = 291. From Eq. 2 in the main text, we calculate *K*_*s, j*_:

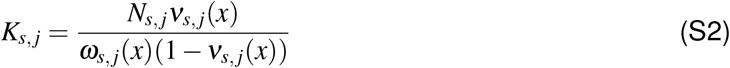

Assuming adapted individuals (*ω*_*s, j*_(*x*) = 1), we can now calculate the carrying capacity parameter 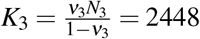. Assuming that *b*_3_ = 64, from Eq. S1 we get the number of stage zero individuals in the littoral environment *N*_0_ = 15125.

We now have to set the other three survival rate *v*_0_, *v*_1_ and *v*_2_. We know their product because it is equal to the ratio of stage 3 and 0 individuals: 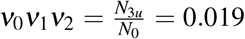. One possible simple combination of surviving rates that appears to be realistic is (*ν*_0_, *ν*_1_, *ν*_2_) = (0.2, 0.25, 0.4). With these values, we have equilibrium densities at (*N*_0_, *N*_1_, *N*_2_, *N*_3*u*_, *N*_3*m*_) = (15125, 3025, 756, 291, 709). Using equation S2, the carrying capacity parameters for sizes 0, 1 and 2 for the littoral morph become *K*_0_ = 3781, *K*_1_ = 1008, *K*_2_ = 504, *K*_3_ = 2448.

**Table S1:** Survival *ν*_*s, j*_ carrying capacity *K*_*s, j*_ at equilibrium for the different stage in different habitats.

| Stage | Littoral | Pelagic | Profundal |
| --- | --- | --- | --- |
| 0 (2cm) | 0.2/3781 | 0.338/2817 | 0.1/73 |
| 1 (15cm) | 0.25/1008 | 0.69/13355 | 0.4/44 |
| 2 (20cm) | 0.4/504 | 0/0 | 0.82/683 |
| 3 (30cm) | 0.71/2448 | 0/0 | 0/0 |

#### Profundal morph

In order to find parameters for the carrying capacity of the produndal population, we need to make further assumptions: First, we set its fertility at *b*_2_ = 16 which fits the known relationship between fish size and the number of eggs produced []schneiderEtAl2000, sandlundEtAl2013. In our unpublished data, of the 243 profundal whitefish sampled, the oldest one was 30 years old. Using the same logic as above, we find its survival rate *ν*_2_ = 0.0.82 at stage two. We set the equilibrium density of the largest profundal fish to *N*_2_ = 150. Using those assumptions we get a profundal niche with the following equilibrium densities (*N*_0_, *N*_1_, *N*_2*u*_, *N*_2*m*_) = (656, 66, 123, 27) and survival (*ν*_0_, *ν*_1_, *ν*_2_) = (0.10, 0.40, 0.82). Using equation S2, the carrying capacity parameters for sizes 0, 1, and 2 for the profundal morph become *K*_0_ = 73, *K*_1_ = 44, *K*_3_ = 683.

#### Pelagic

To extract the survival and carrying capacities of the pelagic population, we again use the relationship between fish fertility and size []schneiderEtAl2000, sandlundEtAl2013 and get *b*_1_ = 4. Of the 913 profundal whitefish sampled, five reached the age of 16 years old. Using the same logic as above, we have a survival rate of *ν*_1_ = 0.0.69 at stage one. In this case, we use *N*_1_ = 6000, and *ν*_1_ = 0.69 and extract the other parameters using the Leslie equations above. We now have the equilibrium densities (*N*_0_, *N*_1*u*_, *N*_1*m*_) = (5517, 1862, 4138) and survival (*ν*_0_, *ν*_1_) = (0.338, 0.69) for the pelagic niche. Using equation S2 results in carrying capacity parameters *K*_0_ = 2817, *K*_1_ = 13355 for the pelagic environment.

Table S1 summarizes the estimates of equilibrium survival rates *v* and *K* parameters.

## Notes

http://xavier.thibert-plante.com/whitefish

